# BET Degraders Reveal BRD4 Disruption of 7SK and P-TEFb is Critical for Effective Reactivation of Latent HIV in CD4+ T-cells

**DOI:** 10.1101/2024.02.23.581756

**Authors:** Anne-Marie W. Turner, Frances M. Bashore, Shane D. Falcinelli, Joshua A. Fox, Alana L. Keller, Anthony D. Fenton, Renee F. Geyer, Brigitte Allard, Jennifer L. Kirchherr, Nancie M. Archin, Lindsey I. James, David M. Margolis

## Abstract

HIV cure strategies that aim to induce viral reactivation for immune clearance leverage latency reversal agents to modulate host pathways which directly or indirectly facilitate viral reactivation. Inhibition of BET (bromo and extra-terminal domain) family member BRD4 reverses HIV latency, but enthusiasm for the use of BET inhibitors in HIV cure studies is tempered by concerns over inhibition of other BET family members and dose-limiting toxicities in oncology trials. Here we evaluated the potential for bivalent chemical degraders targeted to the BET family as alternative latency reversal agents. We observed that despite highly potent and selective BRD4 degradation in primary CD4+ T-cells from ART-suppressed donors, BRD4 degraders failed to induce latency reversal as compared to BET inhibitors. Further, BRD4 degraders failed to mimic previously observed synergistic HIV reactivation between BET inhibitors and an activator of the non-canonical NF-κB pathway. Mechanistic investigation of this discrepancy revealed that latency reversal by BET inhibitors is not related to the abatement of competition between Tat and BRD4 for P-TEFb, but rather the ability of BRD4 to disrupt 7SK and increase the levels of free P-TEFb. This activity is dependent on the shift of BRD4 from chromatin-bound to soluble and retargeting of P-TEFb to chromatin which is dependent on intact BRD4 but independent of the bromodomains.

## Introduction

Advances in antiretroviral therapy have made Human Immunodeficiency Virus (HIV) a chronic but manageable disease. However, a cure remains elusive due to a persistent, transcriptionally repressed latent reservoir in resting CD4+ T-cells (rCD4). One approach to an HIV cure focuses on latency reversal and clearance. A two-component system, latency reversal is dependent on the reactivation of the latent virus reservoir followed by targeted clearance of reactivated cells using methods which boost recognition by the immune system (*1*). Current latency reversal agents (LRAs) rely on disruption or modulation of host pathways which directly or indirectly result in viral reactivation, however, to date, none have proven successful in reducing the size of the latent reservoir (*1*). As such, there is a constant need to identify and evaluate new small molecules targeting well-defined HIV latency pathways for improved specificity and activity. This approach may increase the safety and efficacy of cure studies.

The bromo and extra-terminal domain (BET) family includes ubiquitously expressed BRD2, BRD3, BRD4, and tissue restricted BRDT. All family members contain tandem bromodomains (BD) which bind acetylated substrates, primarily histone tails, and an extra-terminal (ET) domain involved in additional protein-protein interactions (*2, 3*). BRD2/3/4 are recognized global regulators of transcription with BRD4 being the most studied. BRD2 exists as a single protein, BRD3 has a short isoform, BRD3R, which lacks the ET domain, and finally BRD4 has three isoforms, the long isoform BRD4L and two shorter isoforms BRD4Sa and BRD4Sb which differ by a short unique C-terminal sequence on the latter (*2, 3*). BRD4L contains a unique c-terminal domain (CTD) that interacts with positive transcription elongation factor b (P-TEFb) (*2, 3*). P-TEFb, a heterodimer of CyclinT1/T2 and kinase CDK9, is critical for effective release and elongation of paused RNA polymerase II (RNAPII) (*4*). Recruitment of P-TEFb to paused RNAPII by BRD4, the super elongation complex (SEC), or, in the case of HIV, the viral protein Tat, results in CDK9-mediated phosphorylation of the CTD of RNAPII and of repressive factors DSIF and NELF, allowing for productive transcriptional elongation (*4*).

Early studies of the role of BET proteins in HIV focused on the BRD4/P-TEFb axis, demonstrating BRD4 recruitment of P-TEFb could activate reporters containing the HIV promoter (LTR, long terminal repeats) in Tat-free systems (*5*), however overexpression of Tat appeared to compete against this activity (*6*). Later work which specifically identified the terminal 34 amino acids of the BRD4 CTD as critical to the P-TEFb binding also observed that overexpression of a peptide fragment containing amino acids 1209-1362 of BRD4, termed the PID (P-TEFb interacting domain), inhibited Tat-transactivation of the HIV LTR and association with CDK9, suggesting BRD4 and Tat competed for P-TEFb (*7*). Development of one of the first bromodomain inhibitors (BETi) JQ1 in 2010 (*8*) provided the first demonstrations that bromodomain inhibition could reverse HIV latency (*9–12*). These works resulted in the prevailing model that BRD4 bound at the HIV promoter acts to repress HIV transcription by blocking the Tat/P-TEFb interaction and that inhibition and disruption of BRD4 from chromatin by BETi relieves this competition. Two groups further described the ability of JQ1 to disrupt P-TEFb from the repressive non-coding ribonucleoprotein (RNP) complex 7SK, providing an additional mechanism to increase available P-TEFb levels that could contribute to both Tat-dependent and independent activation (*10, 11*). Further work on BRD4 restriction of HIV has implicated the short isoforms of BRD4 (BRD4S), which lack the PID, in recruitment of the repressive BAF complex to maintain the repressive Nuc-1 at the HIV LTR (*13*) and the maintenance of repressive BRD4 at the HIV LTR via KAT5-mediated histone 4 acetylation (H4ac) of the LTR (*14, 15*). Studies using shRNA or CRISPR knockdown of BRD2 suggest some involvement in latency reversal, however whether this is a direct or secondary impact of BRD2 depletion, and the potential mechanisms are currently unknown (*16, 17*).

BRD4 inhibition remains an attractive target for latency reversal, but enthusiasm for the use of BETi in HIV cure studies is tempered by fact that current BETi are pan-inhibitors of all family members and data from oncology studies have demonstrated unfavorable toxicities including thrombocytopenia and severe gastrointestinal symptoms (*18, 19*). A new class of molecules, PROTACs (proteolysis-targeting chimeras), has recently emerged as a potential alternative to BETi. PROTACs, also referred to as bivalent chemical degraders, aim to degrade the protein of interest rather than simply inhibit its function. Specifically, these degraders are a bifunctional molecule that simultaneously binds an E3 ubiquitin ligase and a protein of interest (POI), resulting in ubiquitination of the POI by the E3 ligase complex and POI degradation by the proteasome (Figure 1A). Unique features of chemical degraders include 1) degradation versus inhibition of target molecules, 2) ability to use POI-targeting ligands that do not bind the active site, 3) the catalytic nature of the reagent which allows use at sub-stoichiometric concentrations, and 4) the design of the bivalent molecule can result in improved selectivity over the POI-targeting ligand alone (*20, 21*). Thus, there is a considerable interest in developing degraders for clinical applications (*22*). Further, multiple degraders have been described with BRD4-selective degradation profiles (*23–25*).

**Figure 1–.**
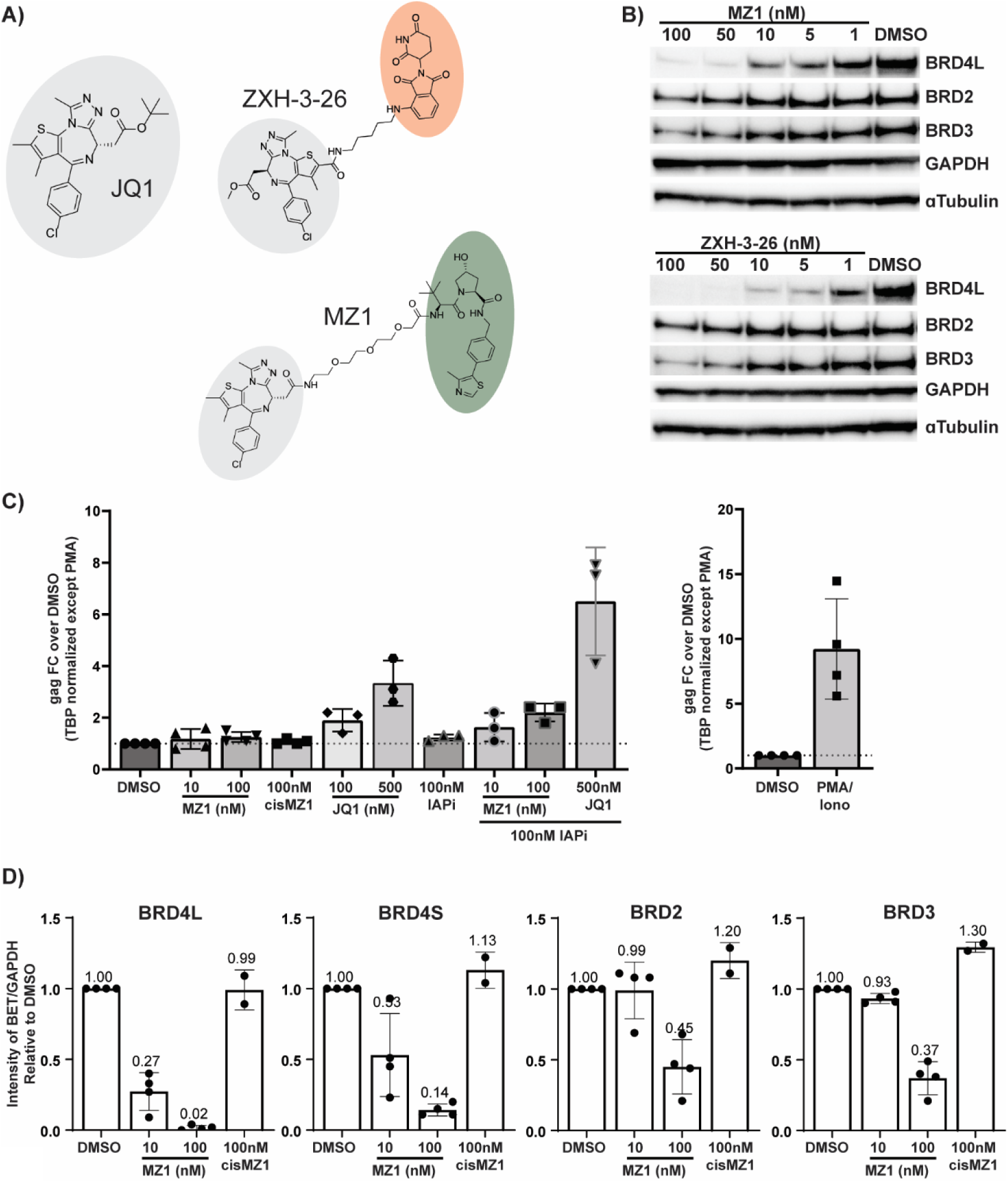
HIV latency reversal in ART-suppressed donors by Bivalent Degraders of the BET proteins –. **(A)** BET inhibitor JQ1 is highlighted in gray. Degraders MZ1 and ZXH-3-26 use JQ1 linked to either VHL (green) or CRBN (orange) recruiting ligands. **(B)** A targeted dose titration of MZ1 and ZXH-3-26 in uninfected primary CD4+ T**-**cells identifies BRD4-specific degradation at 10nM and 5nM respectively Total CD4+ T-cells from ART-suppressed donors were treated for 24 hours with **(C)** MZ1 alone (n=4) or in combination with IAPi AZD5582 (n=3) and induction of gag cell associated RNA assayed by qRT-PCR. **(D)** Cells from MZ1 and control treated cells were assessed by western blot and BET protein levels quantitated relative to GAPDH levels for all donors.

To evaluate the potential for BET degraders as LRAs, multiple compounds were evaluated for target specificity and HIV latency reversal in both Jurkat-derived models and in CD4+ T-cells from ART-suppressed donors. Interestingly, despite the identification of two BET degraders with BRD4 specificity in primary CD4+ T-cells, highly potent BRD4 degradation failed to induce latency reversal in cells from ART-suppressed donors as compared to BETi. Further, BRD4 degraders failed to mimic synergistic HIV reactivation observed between BETi and an activator of the non-canonical NF-κB pathway, as previously described (*16*). We endeavored to understand this discrepancy and found that the primary mechanism of HIV latency reversal by BETi is not the removal of BRD4 from chromatin but the disruption of P-TEFb from repressive the 7SK non-coding RNA complex. Of note, this activity was dependent on the physical BRD4 protein but independent of the bromodomains. These results further emphasize P-TEFb restriction in CD4+ T-cells as a main driver in the maintenance of HIV latency.

## Results

### BRD4-Selective Degradation by MZ1 and ZXH-3-26 in Primary CD4+ T-cells

We selected two commercially available small molecule degraders which used the BETi JQ1 as the targeting moiety linked to ligands recruiting either the Von Hipple-Lindau (VHL) or cereblon (CRBN) E3 ligase complexes that had previously been described as BRD4-selective (Figure 1A). The degrader MZ1 recruits VHL and was described as BRD4 selective below 250 nM (*25*). *cis*MZ1, a diastereomer of MZ1 containing a *cis*-hydroxyproline within the VHL ligand that abrogates VHL binding, was also purchased as a control. ZXH-3-26 recruits CRBN and was described as highly BRD4 selective over a broad range of concentrations (*24*). To determine effective concentrations at which we could reproducibly and selectively degrade BRD4, we treated primary CD4+ T-cells isolated from healthy donors with a broad concentration range for 24 hours and assayed BRD4L/S, BRD2, and BRD3 levels by western (Supplementary Figure S1A-B). We targeted < 100 nM for further testing and observed no impact on protein levels using controls JQ1 and cisMZ1 (Supplementary Figure S1C-D). A targeted dose titration ranging from 1 nM to 100 nM of MZ1 and ZXH-3-26 demonstrated selective degradation of BRD4 at 10 nM by MZ1 and 5nM by ZXH-3-26 (Figure 1B). For both degraders, concentrations ≥50 nM demonstrated some reduction of BRD2 and BRD3 protein levels (Figure 1B).

### Selective BRD4 Degradation Does Not Induce Latency Reversal in HIV+ CD4+ T-cells

We recently published that activation of the non-canonical NF-κB pathway using the Inhibitor of <u>Ap</u>optosis inhibitor (IAPi) small molecule AZD5582 synergizes with BET inhibitors to strongly induce HIV *gag* cell-associated RNA in cells from aviremic donors (*16*). As single LRAs often result in modest latency reversal, we tested ZXH-3-26 and MZ1 alone and in combination with IAPi AZD5582. In Falcinelli et al, we examined the ability of ZXH-3-26 to induce gag cell-associated RNA alone and in combination with AZD5582 (*16*). We observed little to no induction of HIV *gag* cell-associated RNA with ZXH-3-26 alone despite robust BRD4 degradation at 5 nM and no evidence of synergy with AZD5582 (*16*). Here we provide westerns characterizing ZXH-3-26 degradation in 2 donors (Supplementary Figure S2A-B) not previously published and provide calculations of average degradation efficacy for all BET proteins in all 3 donors assayed in Falcinelli et *al* (Supplementary Figure S2C-F), further reinforcing that we were able to achieve robust and selective BRD4 degradation and yet observed no latency reversal.

To extend these observations, we performed matching experiments with MZ1 at 10 nM (BRD4-selective) and 100 nM (pan-degradation). At both concentrations, MZ1 alone failed to induce HIV *gag* cell-associated RNA as compared to JQ1 alone (Figure 1C). Similar to ZXH-3-26, 10 nM MZ1 showed no synergy with AZD5582 as compared to JQ1 (Figure 1C), despite specifically decreasing levels of both the BRD4L and BRD4S isoforms while sparing BRD2 and BRD3 (Figure 1D). Individual western blots for each donor are provided in Supplementary Figure 3A-D. There was minimal impact on cellular viability by MZ1 alone and in combination with AZD5582 via measurement of total cellular ATP levels (Supplementary Figure 3E). Combined with our published ZXH-3-26 data, both BRD4-selective and pan-BET degradation by two potent bivalent degraders failed to reproduce the synergy observed between BETi and IAPi, implicating a non-bromodomain related mechanism in BETi-IAPi synergy for HIV latency reversal in primary CD4+ T cells and requirement of the physical BRD4 protein.

### BRD4-Selective Degradation by BET Degraders Fails to Induce Latency Reversal in Jurkat-derived Latency Models

Knockdown of BRD4 using shRNA/siRNAs approaches have been demonstrated to induce latency reversal in Jurkat-derived models of HIV latency (*11, 17*). We next performed a 16-point dose titration of MZ1 and ZXH-3-26 in the Jurkat-derived JLatA2 reporter line. JLatA2 cells contain an LTR-Tat-IRES-GFP reporter in which LTR activation drives GFP expression (*26*) and have previously been used in the study of BRD4 in HIV latency (*11, 17*). JQ1(+) and stereoisomer negative control JQ1(-) (*8*) were used to compare degrader LRA activity to the parent BETi molecule. Latency reversal by JQ1(+) is graphed in the open circles in all graphs to allow direct comparison. As expected, JQ1(+) induced latency reversal as measured by percent GFP positive JLatA2 cells by flow cytometry, with maximal activation inducing approximately 30% of the population (Figure 2A and D, open circles). JQ1(-) showed no latency reversal (Supplementary Figure 4A, open squares). Neither compound showed a measurable impact on cellular viability as measured by a fixable live/dead stain (Supplementary Figure 4A, dark gray filled symbols). JQ1(+) treatment did not induce any changes in protein level of any BET family members (Supplementary Figure 4B-C).

**Figure 2–.**
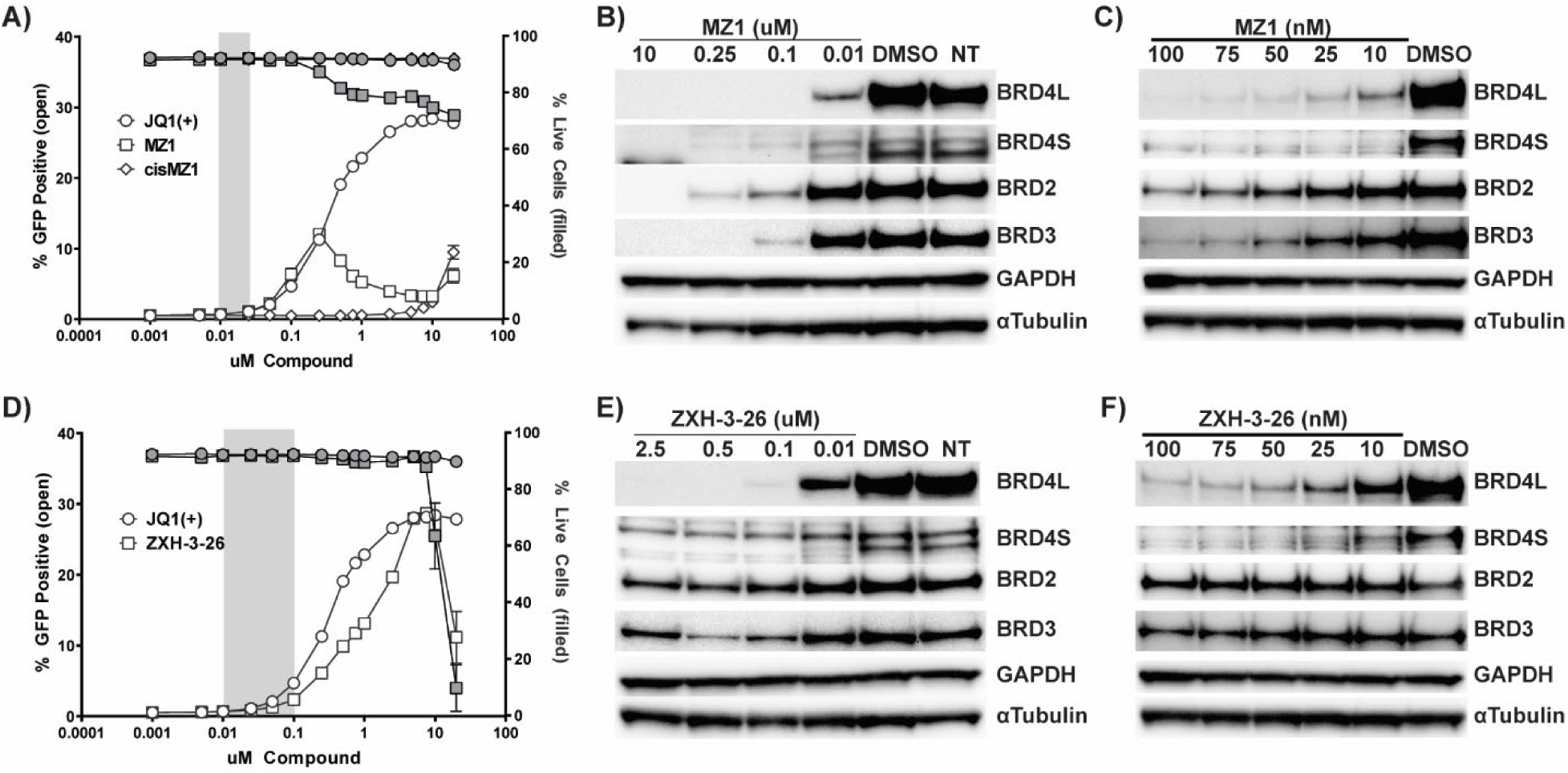
Latency reversal and targeted degradation of BET degraders in a Jurkat-derived Latency Model –. Latency reversal was assessed via treatment of JLatA2 cells with a 16-point dose titration of **(A)** MZ1 and *cis*MZ1. JQ1(+) data is overlayed and provided in Supplementary Figure SI3. Each titration was performed 3 independent times with triplicate treatments for each experiment (n=9) with GFP (open symbols) assessed by flow cytometry as a measure of latency reactivation and viability by live/dead stain (gray symbols). The window of BRD4 specific degradation for MZ1 is highlighted in gray as determined by corresponding westerns for both BRD4L(ong) and BRD4S(hort) isoforms as well as BRD2 and BRD3 in JLatA2 cells **(B-C)**. Matching experiments were performed for **(D)** ZXH-3-26 and the window of BRD4-selective degradation is highlighted in gray based on corresponding westerns **(E-F)**.

While both MZ1 and ZXH-3-26 demonstrated latency reversal in JLatA2 cells as measured by GFP induction (Figure 2A and D), this occurred outside the range of BRD4-selective degradation as assayed by western blot using both extended and targeted concentrations for both degraders (Figure 2B, C, E, F). The BRD4-selective range for both degraders is highlighted in gray in Figures 2A and 2D. MZ1 displayed a pattern whereby GFP levels initially increased but then reached a concentration at which both GFP and cellular viability began to decrease (Figure 2A). The concentrations at which this occurred correlated with near complete degradation of all BET family members as measured by western blot (Figure 2B). These observations are consistent with previously published work by Winters *et al*. which observed significant defects in transcriptional initiation and elongation when using the BET degrader dBET6 at pan-degrading concentrations (*27*).

We observed some indication of hook effect, the concentration at which the bivalent molecule is saturating such that the ternary complex is not formed and the individual parent inhibitors can act independently (*20, 21*), by both degraders. Both MZ1 (Figure 2A, open squares) and *cis*MZ1 (Figure 2A, open diamonds) at concentrations ≥ 10 µM induced GFP expression in JLatA2 cells. Poor cellular permeability due to the high molecular weight of chemical degraders has been characterized, with a recent study demonstrating MZ1 is over 165,000-fold less permeable than JQ1 (*28*). GFP induction by *cis*MZ1 at 10 µM suggests the intracellular concentration of *cis*MZ1 approaches concentrations at which the JQ1 moiety alone can induce latency reversal. However, we did not observe a return of protein levels at 10 µM by western at the 24 hour timepoint which would have indicated hook effect. Similar to primary CD4+ T-cells, ZXH-3-26 was the most potent and selective in JLatA2 cells, as we never observed a complete loss of BRD2 or BRD3 (Figure 2E-F). We did observe recovery of BRD2 and BRD3 proteins levels in ZXH-3-26 treated cells at 2.5 µM, indicating hook effect, but BRD4 protein levels did not show recovery (Figure 2E). Coupled with our observations of potential hook effect as measured by GFP induction by MZ1/*cis*MZ1 but no observable protein recovery by western, it may be that the recovery of BRD4 is either under our limit of detection by western or BRD4 recovery differs from BRD2/3 depending on selectivity of the degrader. However, for both degraders this occurred at significantly higher concentrations than those at which BRD4-selective degradation occurred and at which we assayed for latency reversal and thus had no relevance to the lack of observed induction of HIV.

### Viral Reporter Contributes to Perceived Latency Reversal and IAPi Synergy in Jurkats

While treatment of JLatA2 cells with MZ1 and ZXH-3-26 did not demonstrate latency reversal at BRD4 selective concentrations, we observed induction of GFP at higher, non-selective concentrations (Figure 2). We sought to determine if this was unique to the JLatA2 line as different reporter lines are known to demonstrated variable latency reversal in response to the same LRA (*29*). As previously described, JLatA2 cells contain a short, 1.5kB reporter. JLat10.6 cells contain a defective but full-length viral reporter containing a frameshift in *env* and GFP in the place of *nef* (*26, 30*). When treated with the cytokine tumor necrosis factor alpha (TNFα), JLat10.6 cells reactivate to similar levels as JLatA2 cells (Supplementary Figure 5A-B). We performed the same 16-point titration of MZ1, *cis*MZ1, ZXH-3-26, and JQ1 in JLat10.6 cells. While JQ1 latency reversal was also attenuated as compared to JLatA2 cells, both ZXH-26 and MZ1 failed to induce notable latency reversal in JLat10.6 cells (Figure 3A and Supplementary Figure 5C respectively).

**Figure 3–.**
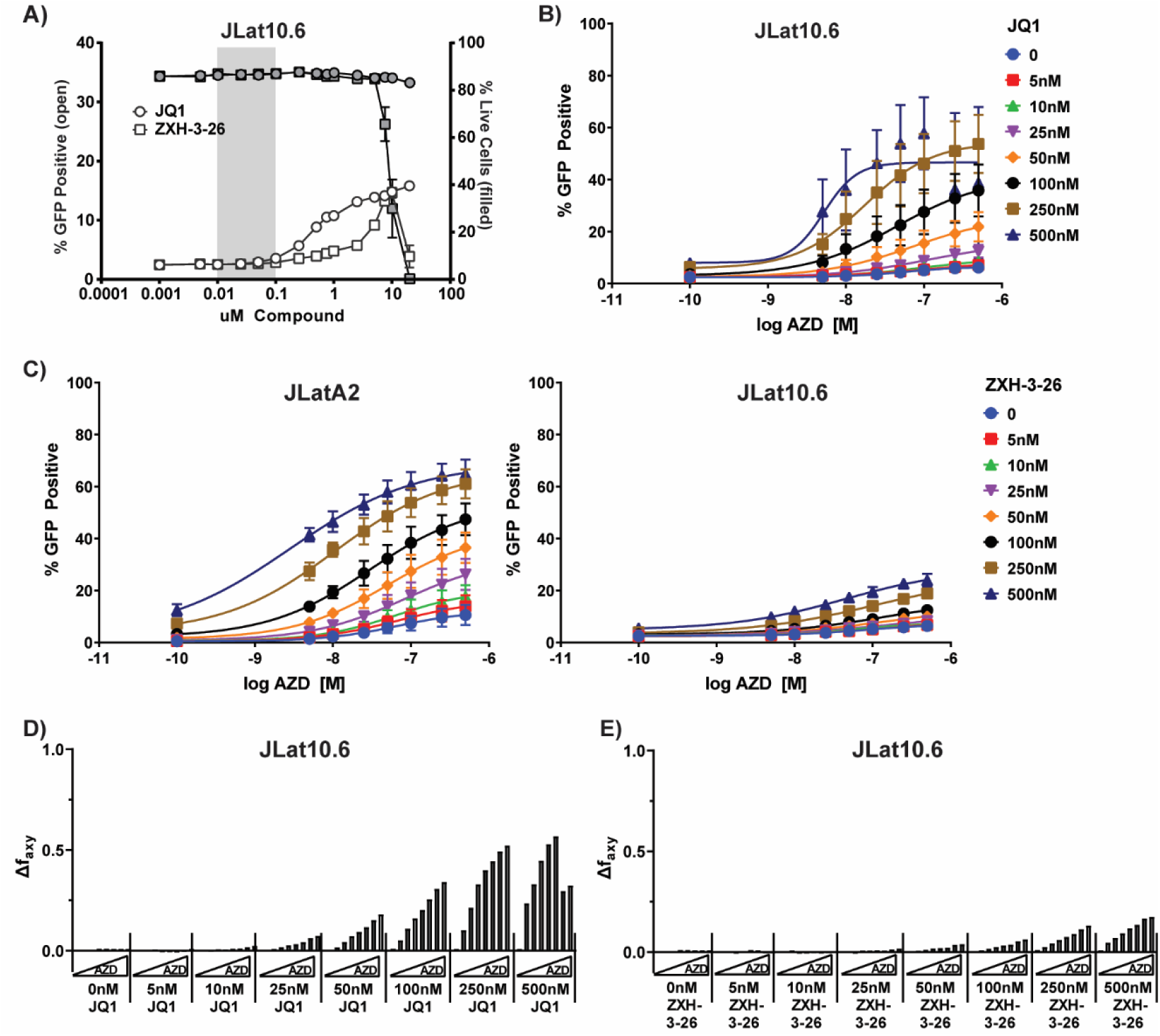
Latency reversal and synergy with AZD5582 by degrader ZXH-3-26 is attenuated in a full length Jurkat cell line –. **(A)** 16-point dose titration ZXH-3-26 and JQ1 in JLat10.6 cells demonstrates minimal latency reversal by ZXH-3-26 and reduced activity by JQ1(+) compared to JLatA2 despite similar maximal activation by TNFα (SI4A-B). **(B)** An 8-point dose titration of AZD-5582 and JQ1 demonstrates synergy in JLat10.6 cells. **(C)** AZD-5582 and ZXH-3-26 demonstrate synergy in JLatA2 cells but demonstrate no synergy at BRD4-selective doses of ZXH-3-26 in JLat10.6 cells **(C)**. Bliss synergy calculations of AZD5582/JQ1 **(D)** and AZD5582/ZXH-3-26 **(E)** in JLat10.6 cells. Latency reversal by MZ1 in JLat10.6 cells, synergy experiment in JLatA2 cells for AZD5582/JQ1, and additional Bliss synergy graphs are provided in Supplementary Figure 4.

We next examined the impact of the reporter line on our observations of synergy between BETi, BET degraders, and IAPi. We performed an 8-point cross titration between AZD5582 and either JQ1 or ZXH-3-26 in both cell lines and used the Bliss Independence Model to analyze whether the combinations displayed antagonism (Δf_axy_<0), independence (Δf_axy_=0), or synergy (Δf_axy_>0). Consistent with our previous observations, AZD5582 in combination with JQ1 induced synergy in both cell lines (Figure 3B and Supplementary Figure 5D). Interestingly, ZXH-3-26 in combination with AZD5582 in JLatA2 cells resulted in reduced but observable synergy even at BRD4-selective concentrations (< 100 nM) (Figure 3C) while little to no synergy was observable at the same concentrations in JLat10.6 cells (Figure 3D). Calculations of Bliss synergy for cross titrations are provided in Figures 3D-E for the JLat10.6 line and Supplementary Figure 5E-F for the JLatA2 line.

The lack of observable synergy in JLat10.6 cells was consistent with our prior results in Falcinelli et al (*16*) using another full-length HIV reporter cell line model. We hypothesize the smaller reporter virus construct in the JLatA2 reporter line has a lower barrier to productive transcriptional elongation and is more susceptible to secondary cellular effects of BRD4 loss which may occur rapidly at higher degrader concentrations. Taken together with the observation that BRD4-selective concentrations fail to induce latency reversal in a broad dose-response curve, we find rapid and robust BRD4 degradation by small molecule degraders does not induce HIV latency reversal in primary cells or Jurkat reporter models.

### Modulation of P-TEFb/7SK Differs Between BRD4 Degradation and Inhibition

Three proposed mechanisms for the latency reversal activity of BETi are (1) reduction of competition for P-TEFb between Tat and BRD4, (2) release of P-TEFb from the 7SK RNP, and (3) maintenance of repressive BAF at the LTR by BRD4S (*9–13*). We demonstrate both MZ1 and ZXH-3-26 degrade the BRD4S isoform with no impact on latency reversal (Figures 1 and 2). To further understand how BETi but not BET degraders reverse HIV latency, we performed immunoprecipitations of CyclinT1 to identify levels of interacting BRD4 to assess competition for available P-TEFb and examined HEXIM upregulation as a proxy for P-TEFb release. The 7SK RNP is composed of the 331 nucleotide non-coding RNA 7SK and major proteins LARP7, MeCBP, and HEXIM (*4*). HEXIM (Hexamethylene-bis-acetamide Inducible Transcript 1) directly interacts with stem loop 1 (SL1) of 7SK and with the P-TEFb heterodimer, blocking the ATP binding pocket of CDK9 and sequestering P-TEFb in this repressive RNP complex (*31, 32*). Disruptions of the 7SK/P-TEFb equilibrium by cellular stress, stimuli, or small molecules have been previously shown to induce a feedback loop in which release of excess P-TEFb triggers an upregulation of HEXIM1 gene transcription and protein levels, resulting in re-sequestration of active P-TEFb (*4, 33, 34*).

As expected, CyclinT1 co-immunoprecipitated with BRD4, CDK9, HEXIM, and SEC member AFF4 but not BRD2, a non-interacting control (Figure 4A). JQ1 treatment resulted in increased BRD4 association with CyclinT1 at 500nM (Figure 4A) and modest increase at 250nM (Supplementary Figure 6A). These results demonstrate BETi do not impact BRD4/P-TEFb interaction, consistent with the BRD4 CTD/PID and not the bromodomains as the primary interaction domain with CyclinT1 but inconsistent with a mechanism in which BRD4 inhibition reduces competition for P-TEFb with Tat. Meanwhile, ZXH-3-26 treatment abrogated BRD4 interaction with CyclinT1 (Figure 4A), suggesting BET degraders would effectively reduce competition between BRD4 and Tat for P-TEFb if this were the primary mechanism of latency reversal. These results combined with our observations that robust BRD4 degradation fails to induce latency reversal suggests that the prior interpretation that BRD4 is repressive and removal from chromatin dictates latency reversal merits reevaluation.

**Figure 4–.**
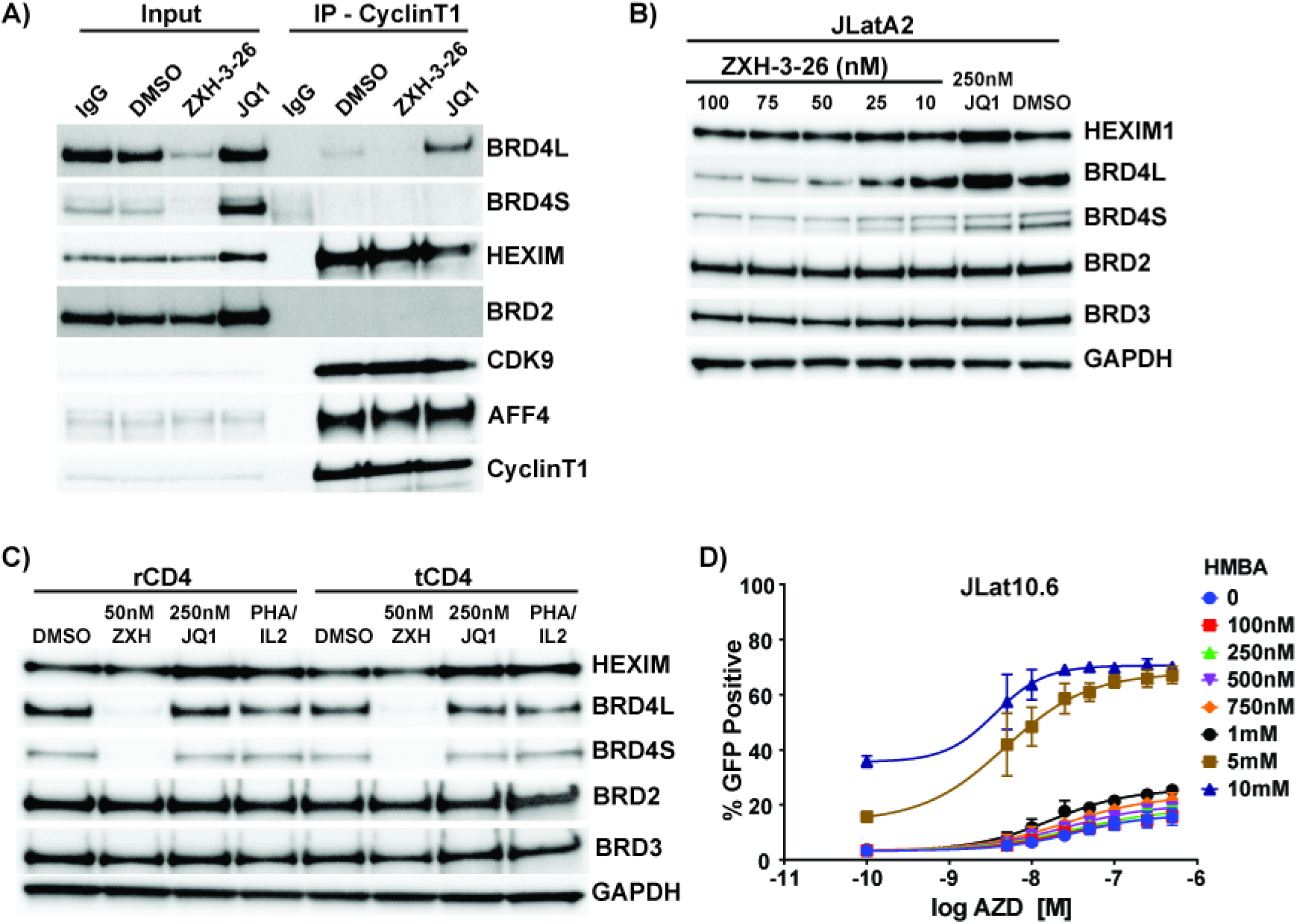
HEXIM1 is upregulated in response to BETi but not BET PROTACs –. **(A)** Jurkat cells were treated with vehicle control (DMSO), 50nM ZXH-3-26, or 500nM JQ1 for 24hrs followed by a CyclinT1 or IgG immunoprecipitation and western blot for associated proteins. HEXIM1 protein levels are upregulated in **(B)** Jurkat-derived cells in response to JQ1 but not in response to ZXH-3-26. This observation is repeated in both **(C)** resting and total primary CD4+ T-cells. **(D)** Latency reversal in JLat10.6 cells with an 8-point dose titration of AZD5582 and HMBA. Data represents 3 independent experiments of each 8-point titration.

Interestingly, immunoprecipitation input samples demonstrated JQ1 strongly induced HEXIM expression in contrast to ZXH-3-26 (Figure 4A). Over the optimized dose range in JLatA2 cells, ZXH-3-26 at concentrations selective for BRD4 degradation did not induce increased HEXIM protein levels as compared to JQ1 (Figure 4B). We observed similar results with MZ1 (Supplementary Figure 6B) at 10-100 nM and for both degraders at a greater range of concentrations while JQ1 strongly induced HEXIM at concentrations as low as 100 nM after 24 hours (Supplementary Figure 6C). Importantly, we confirmed HEXIM upregulation by BETi but not BET degraders in both total and, importantly, transcriptionally quiescent resting CD4+ T-cells (CD25/CD69/HLA-DR negative) (Figure 4C). While HEXIM upregulation is an indirect measure of P-TEFb release, these results strongly support the hypothesis that the primary difference between BET degraders and BETi is the disruption of P-TEFb from the 7SK RNP and implicates this as the primary mechanism in HIV latency reversal.

To further confirm BETi/IAPi synergy is primarily driven by P-TEFb disruption from 7SK, we hypothesized pairing AZD5582 with another disruptor of the P-TEFb pathway, hexamethylene bisacetamide (HMBA), would also result in synergistic latency reversal. Early studies of HMBA observed the upregulation of HEXIM mRNA as a primary result of treatment (*35*), an observation now understood to be a result P-TEFb disruption (*34, 36, 37*). HMBA is also a known LRA requiring high mM concentrations for activity (*38, 39*). When combined with AZD5582, we observed weak synergy at concentrations ≤ 1 mM but observed strong synergy at 5 mM and 10 mM with suboptimal concentrations of AZD5582 in JLat10.6 cells (Figure 4D and Supplementary Figure S6D), confirming P-TEFb release from 7SK as the primary driver of synergy with IAPi.

### Disruption of P-TEFb/7SK is Dependent on Release of Intact BRD4 from Chromatin

The specific mechanism through which BETi disrupt P-TEFb from 7SK has not been described. Induction of the HEXIM transcript, which appears to mirror HIV activation in this context, is reported to be a direct feedback loop dependent on P-TEFb and SEC components but independent of BRD4 (*33, 40*). To further probe the mechanism of JQ1-mediated P-TEFb release, we decided to use HEXIM1 upregulation as a reporter and with some modification, replicated work published by Lui et al (*33*), in which they identified the minimal HEXIM promoter responsive to modulation of P-TEFb levels. We cloned the 104bp of the HEXIM1 promoter identified by Liu et al but also included the entire HEXIM1 untranslated region (UTR) upstream of luciferase and generated 293T/17 cell lines which stably expressed the minimal promoter construct (293T-HEXIMpr104) and a control reporter that only contained the UTR (293T-HEXIMpr0) (Figure 5A). Preliminary characterization confirmed only the 293T-HEXIMpr104 line was responsive to JQ1 and HMBA while Flavopiridol, a CDK9 inhibitor, reduced basal luciferase expression (Supplementary Figure 7A-B). Consistent with our western results and Lui et al (*33*), we observed various concentrations of JQ1 induced luciferase expression (3.32X-4.56X induction over control, Figure 5B), demonstrating that HEXIM protein induction occurs secondary to transcriptional upregulation and is not a result of stabilization of the protein. We next examined the impact of BET degraders on HEXIM1 transcriptional induction. ZXH-3-26 at a BRD4-selective dose of 50 nM failed to induce a response (Figure 5B). At higher, non-BRD4 selective doses of 100 and 250 nM, ZXH-3-26 treatment resulted in a mild upregulation (1.2-1.36X over control) but remained weaker than HMBA (2.58X induction) and JQ1 (3.23-4.56X induction over control) (Figure 5B). Taken together, these data suggest that BETi but not BET degraders mediate P-TEFb disruption from 7SK RNP (as measured by HEXIM1 upregulation), and this disruption correlates with HIV latency reversal.

**Figure 5–.**
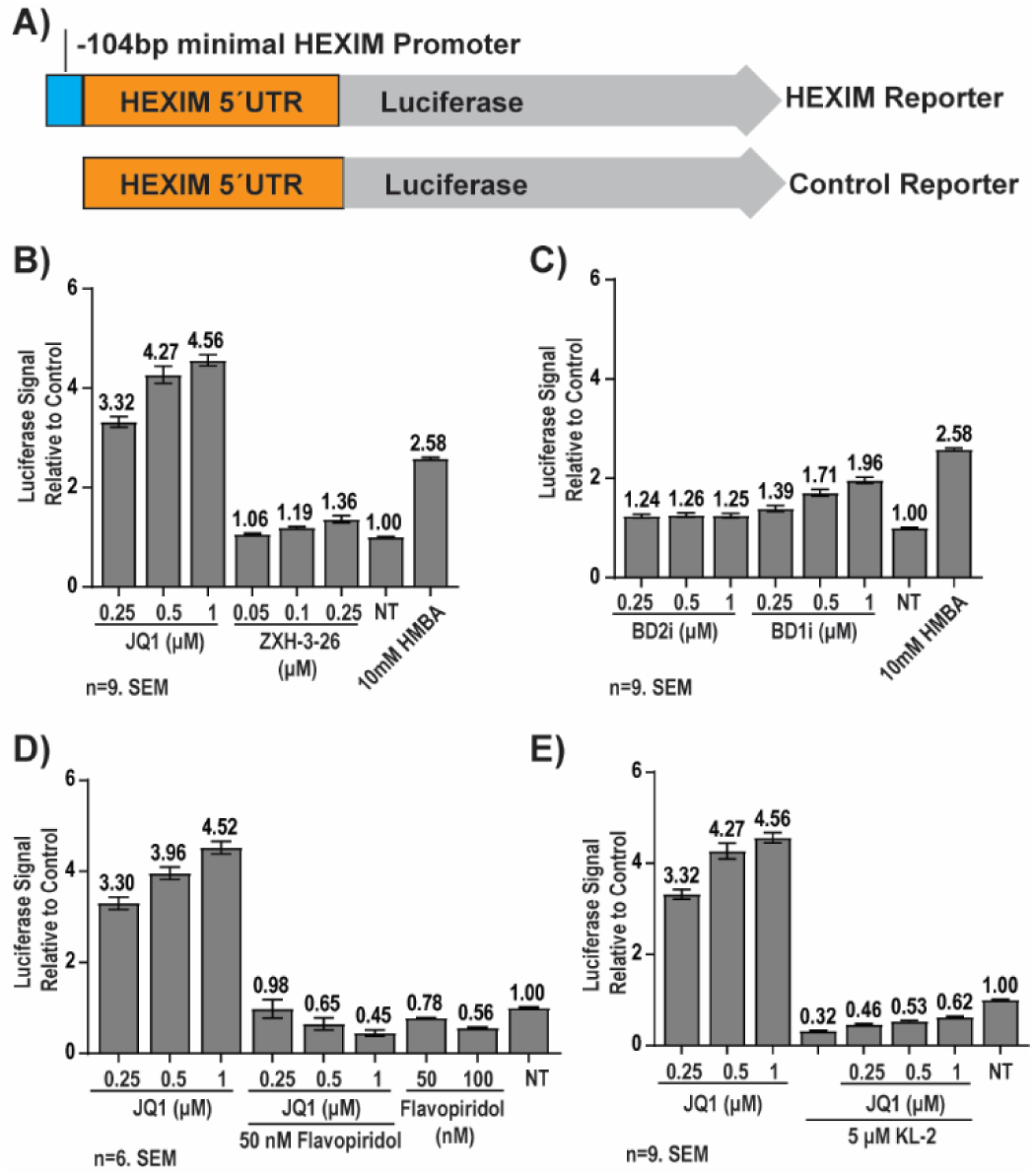
HEXIM1 induction occurs at the transcriptional level and requires the SEC –. To generate 293T cell lines for high throughput assay of conditions which induce the HEXIM promoter, **(A)** the reported minimal HEXIM1 promoter and UTR or the UTR alone (control) were inserted before a luciferase reporter. Fold induction of luciferase is demonstrated for **(B)** JQ1 and ZXH-3-26 and for **(C)** BD1 or BD2 specific inhibitors. Cells were treated at indicated concentrations for 24 hrs and luciferase signal standardized to no treatment (NT, DMSO control). Luciferase induction by JQ1 was blocked by concurrent treatment with **(D)** Flavopiridol and **(E)** KL-2.

Next, we evaluated whether displacement of BRD4 from chromatin is necessary for P-TEFb disruption and subsequent HEXIM upregulation. The BET family contains tandem bromodomains, referred to as BD1 and BD2. We next used recently published BD-specific inhibitors to determine the contribution of each bromodomain to HEXIM induction. BD2 inhibitor GSK-046A (*41, 42*), which does not disrupt BET protein binding to chromatin, weakly upregulated HEXIM (1.25X over the control) but showed no dose-dependent increase in luciferase expression (Figure 5C). In contrast, BD1 inhibitor GSK-779A (*41, 42*), which results in disruption of BET proteins from chromatin similar to BETi, resulted in a dose-dependent increase in luciferase expression (Figure 5C). These results indicate the release of BRD4 from chromatin is necessary for P-TEFb disruption. We further assayed the ability of CDK9 inhibitor Flavopiradol and compound KL-2 (*43*), which inhibits the AFF4-P-TEFb SEC interaction, to block HEXIM upregulation. While both compounds exhibited some level of toxicity (Supplementary Figure 6E-F), both blocked induction of luciferase by JQ1 (Figure 5D-E), confirming P-TEFb and AFF4 have roles in HEXIM upregulation (*33*).

### BRD4 Is Required for Upregulation of HEXIM Transcription

These data implicate BRD4 release from chromatin as critical for P-TEFb disruption by BETi, however degradation of BRD4 abrogates HEXIM1 upregulation and associated latency reversal activity. Collectively, this suggests that while bromodomain inhibition and subsequent chromatin displacement of BRD4 are important, other domains of BRD4 play a critical role in the actual disruption of P-TEFb from 7SK RNA, HEXIM upregulation, and latency reversal. In addition to the well characterize CTD/PID (*7, 44*), the BD domains, specifically BD2, have also been implicated in binding acetylated CyclinT1 and necessary for latency reversal (*45*), however the role of these domains in HEXIM upregulation is unknown. Therefore, we next determined the contribution of the BD-domains and CTD to HEXIM protein induction by overexpressing wild-type BRD4 (BRD4-F-WT), a mutant with non-functional BD domains (BRD4-F-ΔBD), a mutant with the CTD/PID deleted (BRD4-F-ΔCTD), a dual BD/CTD mutant (BRD4-F-ΔBDΔCTD), and a V5-tagged BRD2 and GFP overexpression constructs as controls (Figure 8A). All mutant constructs expressed with only the BRD4-F-ΔBDΔCTD showing reduced proteins levels when assayed by western blot (Figure 6A). Of the expressed proteins, only BRD4-WT and BRD4-F-ΔBD were capable of strongly inducing HEXIM expression (Figure 6A), demonstrating the BD domains are not necessary for this activity. BRD4-F-ΔCTD showed a mild upregulation, however since this construct has intact BD-domains, it is possible that it could displace endogenous WT BRD4 from chromatin, resulting some disruption of P-TEFb. Previously, overexpression of the CTD alone (PID aa1209-1362) was shown to disrupt P-TEFb from HEXIM and inhibit Tat-based elongation of HIV (*7, 44, 45*). Interestingly, we observed overexpression of the CTD/PID alone did not induce HEXIM protein expression (Figure 6B).

**Figure 6–.**
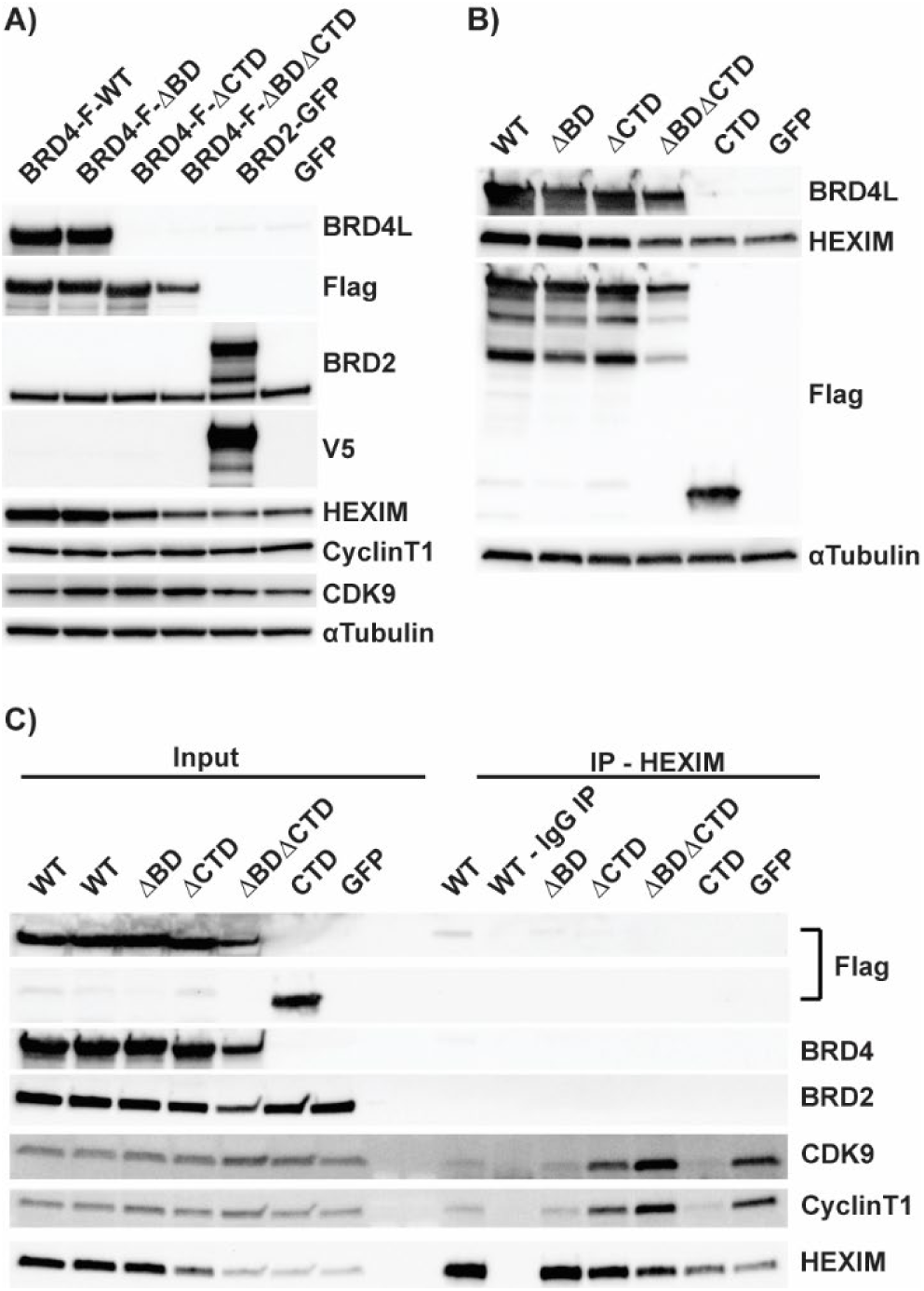
P-TEFb disruption from 7SK is dependent on the CTD but is independent of the BD domains –. **(A)** Overexpression plasmids containing various full-length flag-tagged BRD4 constructs or **(B)** full length and CTD-only constructs were transfected into 293T cells for 48hrs to determine impact on HEXIM1 protein levels. **(C)** Overexpression constructs containing full length WT, mutant or the CTD domain alone were transfected into 293T cells for 48hrs and followed by a HEXIM1 or control IgG immunoprecipitation and western blot for associated proteins.

We then performed HEXIM immunoprecipitations to determine which constructs could result in loss of P-TEFb/HEXIM association in the context of the 7SK RNP to determine how this correlated to HEXIM protein induction. BRD4-WT, BRD4-F-ΔBD, and the CTD/PID alone resulted in a loss of both CyclinT1 and CDK9 association with HEXIM (Figure 6C), irrespective of induced HEXIM protein levels as observed in the input samples. Independent replicates of all blots and immunoprecipitations are provided in Supplementary Figure 8. These results confirm the CTD/PID is required and sufficient for disruption of P-TEFb from the 7SK RNP, but further find HEXIM induction at the transcriptional level is also dependent on intact BRD4 protein and cannot be induced by overexpression of the CTD/PID alone. These data suggest additional protein-protein interactions, potentially mediated through the ET domain or other regions of BRD4, are required to induce HEXIM upregulation and may also be required for HIV latency reversal.

### Latency Reversal by BETi in Jurkats is Dependent on CDK9 but not the SEC

Finally, we moved back into both the JLatA2 and JLat10.6 latency models to determine if the same determinants of HEXIM induction correlated to latency reversal. As expected, Flavopiridol inhibited latency reversal induced by JQ1 in both cell lines (Figure 7A and 7B, viability provided in Supplementary Figure 8A-B). Unexpectedly, SEC inhibitor KL-2 enhanced JQ1-mediated latency reversal (Figure 7A and 7B). Thus, while HEXIM1 transcriptional induction is dependent on the SEC, these results demonstrate JQ1-induced latency reversal of HIV is independent of the SEC and that disruption of other canonical P-TEFb partners beyond the 7SK RNA could potentially feed into the pathway that benefits viral activation in JLat models.

**Figure 7–.**
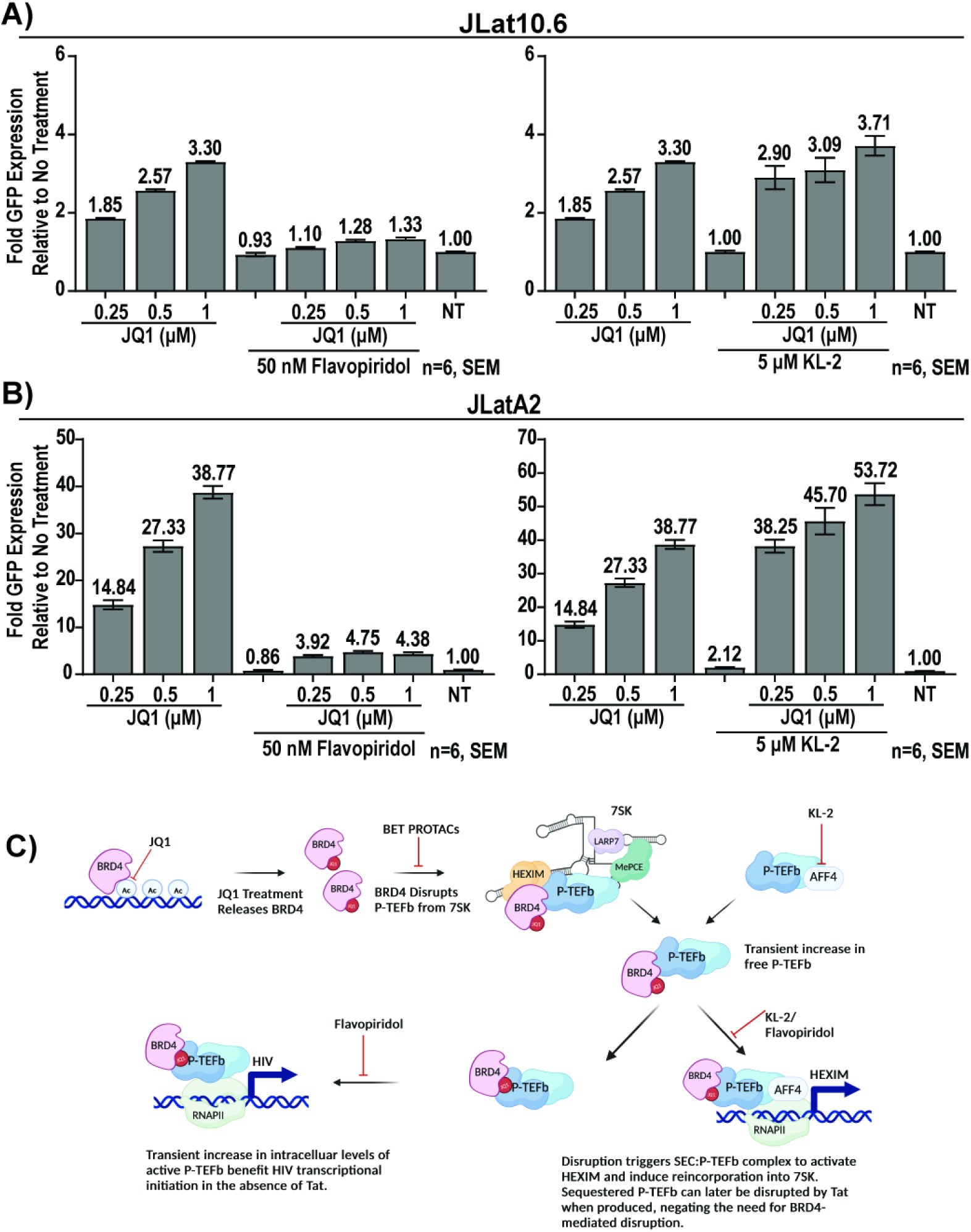
Latency Reversal by JQ1 is blocked by Flavopiridol but not KL-2. In both **(A)** JLat10.6 and **(B)** JLatA2 cells, latency reversal as measured by fold GFP induction by JQ1 is blocked by CDK9 inhibitor but enhanced by SEC inhibitor KL-2. Data represents 3 biological replicates performed during 2 independent experiments (n=6). **(C)** Proposed mechanism of BETi-mediated latency reversal in the absence of Tat.

## Discussion

Here we demonstrate that the mechanism of latency reversal by BETi is not related to the abatement of competition between Tat and BRD4 for P-TEFb but rather the ability of BRD4 to disrupt and increase the levels of free P-TEFb in the cell. Two bivalent small molecule degraders tested here, ZXH-3-26 and MZ1, failed to induce latency reactivation in full-length JLat models and in cells from ART-suppressed donors despite highly selective and robust BRD4 degradation, in direct conflict with a model in which BRD4 at the HIV LTR acts to repress and compete against Tat for P-TEFb. Rather, we observed the primary difference between BRD4 degradation versus inhibition was related to the disruption of P-TEFb from the repressive 7SK complex by BRD4. We propose an updated model, in which release of BRD4 from chromatin by BD-domain inhibition allows BRD4 to interact via the CTD/PID with P-TEFb in 7SK, transiently increasing free P-TEFb levels (Figure 7C). This disruption of P-TEFb equilibrium activates a CDK9 and SEC-dependent feedback loop that triggers an upregulation of the repressive protein HEXIM to reincorporate and re-repress dysregulated P-TEFb. Concurrently, the disruption and increased P-TEFb also triggers HIV LTR activation that is CDK9 dependent, but, in the context of Jurkat-derived models containing Tat, is independent of the SEC (Figure 7C). This work and model further demonstrate that BRD4-selective degraders are not viable alternatives for BET inhibitors as latency reversal agents in the HIV cure space.

This mechanism, however, reconciles observations from numerous studies in the HIV field related to BRD4 and latency reversal. We find that while the CTD/PID can indeed disrupt P-TEFb from 7SK, it cannot activate HEXIM transcription, suggesting targeting of newly released P-TEFb back to chromatin and, by extension, HIV, is dependent on full-length BRD4 but not the bromodomains or histone binding. In support of this, a recent study observed an increased association of CDK9 with active transcriptional start sites and enhancers upon JQ1 treatment via chromatin immunoprecipitation (ChIP)-seq, even in the context of BRD4 loss from chromatin but not upon treatment with degrader dBET6 (*27*). Retargeting of P-TEFb to chromatin likely involves protein-protein interactions mediated by another BRD4 domain that may be beneficial to HIV reactivation in the absence of Tat.

Upon production of Tat, we propose the dynamics of P-TEFb recruitment change. Tat has been shown to disrupt P-TEFb from the 7SK RNP via the arginine-rich RNA binding domain which recognizes the same region of the 7SK RNA as HEXIM, SL1, allowing Tat to directly disrupt and recruit P-TEFb from 7SK (*44, 46*). However, Bisgrove et al demonstrated Tat cannot displace P-TEFb from the BRD4/P-TEFb complex (*7*). We would postulate overexpression of BRD4 or the CTD/PID alone sequesters P-TEFb into a complex which Tat cannot disrupt. Further, lack of HEXIM induction by CTD/PID overexpression would prevent re-sequestration of P-TEFb back into the 7SK RNP, depleting this source of P-TEFb for Tat. Prolonged overexpression of full length BRD4 could also result in a chronic depletion of P-TEFb from 7SK, regardless of HEXIM activation, resulting in the observations that BRD4 and/or CTD/PID overexpression is refractory to Tat-mediated HIV activation (*6, 7, 45*). In the context of BETi-mediate latency reversal, the increased BRD4/P-TEFb complex may result in sufficient HIV transcriptional activation to drive Tat protein expression, at which point eventual loss of the small molecule combined with HEXIM upregulation restores the 7SK/P-TEFb pool, allowing Tat to access P-TEFb and further drive HIV transcription. BRD4 is also reported to have intrinsic kinase activity and can phosphorylate the RNAPII CTD and CDK9 (*47, 48*), activity which may further benefit HIV activation. The mechanisms of BRD4-mediated P-TEFb retargeting and transcriptional activation in the context of bromodomain inhibition merits additional investigation to understand the BRD4/P-TEFb axis and HIV reactivation.

Multiple groups have also linked histone acetyltransferase complexes to the maintenance BRD4 at the HIV LTR resulting in enforcement of latency (*14, 15*). Based on our results, we would hypothesize that targeting complexes which reduce global histone acetylation may also induce release of BRD4 from chromatin in a manner similar to BETi and result in P-TEFb disruption. Related to this, two mechanisms for HMBA disruption of P-TEFb have been proposed. While one links HMBA activation of the PI3K/Akt pathway and the phosphorylation of HEXIM (*37*), another links HMBA treatment to a loss of BRD4 due to a reduction in H4 acetylation and demonstrates an increased association of P-TEFb with non-chromatin bound BRD4 (*36*). Here the authors propose a similar mechanism to us, in which cellular perturbations and changes in histone acetylation can trigger BRD4 release from chromatin to allow P-TEFb recruitment and a switch of BRD4 from chromatin targeting to promoting transcriptional regulation (*36*), suggesting cellular signals which shift histone acetylation may indeed promote HIV reactivation via changes in P-TEFb levels.

Most importantly, we find that the mechanism of P-TEFb disruption from 7SK RNA is active in primary CD4+ T-cells and that HEXIM protein expression can be induced in resting cells. Therefore, despite prior observations of highly restricted CyclinT1 levels in resting cells (*49–51*), we find there is a pool of 7SK-bound P-TEFb that can be disrupted with BET inhibitors but not BET degraders, and results in transcriptional activation. This observation supports P-TEFb disruption from the 7SK RNP as the primary mechanism of BETi-mediated induction of HIV cell-associated RNA in cells from ART-suppressed donors where Tat is presumed to be limited or not present. Future studies focused on targeted ways to specifically disrupt this pool and to identify the specific mechanism and interaction partners involved in BRD4-mediated P-TEFb recruitment and elongation may reveal pathways which can be exploited for HIV cure strategies.

### Methods and Materials

#### Cell Lines

JLatA2 and JLat10.6 (*26, 30*) were obtained from the NIH AIDS Reagent Program. Cells were maintained in RPMI1640 (LifeTech) supplemented with 10% FBS (Millipore) and 100U/mL Pen/Strep (LifeTech) at 37°C/5%CO_2_. HEK293T/17 cells were obtained from ATCC (CRL-11268) and maintained in DMEM (LifeTech) supplemented with 10% FBS (Millipore) and 100U/mL Pen/Strep (LifeTech) at 37°C/5%CO_2_.

#### CD4 T-Cell Isolation

Standard isolation of peripheral blood mononuclear cells via Ficoll-Paque (GE Lifesciences) were performed on samples from anonymous, healthy blood donors (New York Blood Center). Total or resting CD4+ cells were obtained using either the EasySep Human CD4+ T Cell Isolation or EasySep Human Resting CD4+ T-cell Isolation kits (Stemcell Technologies) per manufacturers protocol.

#### Latency Reversal Agents

Compounds were dissolved in DMSO unless indicated. For high-throughput flow and synergy assays, an HP D300e digital dispenser using T8+ or D8+ dispense heads was used to plate DMSO-soluble compounds. ZXH-3-26 (6713, CAS 2243076-67-5) was obtained from Tocris Bioscience. ARV771 (HY-100972, CAS: 1949837-12-0), MZ1 (HY-107425, CAS: 1797406-69-9), *cis*MZ1 (HY-107425A, CAS: 1797406-72-4), dBET1 (HY-101838, CAS: 1799711-21-9), and KL-2 (HY-123972, CAS: 900308-51-2) were purchased from MedChemExpress LLC. HMBA (aqueous) (H4663, CAS: 564468-51-5) was purchased from Millipore Sigma. Flavopiridol (L86-8275, CAS: 146426-40-6) was purchased from Selleckchem. (+)-JQ-1 was purchased from AstaTech Inc. (Catalog #: 41223, CAS: 1268524-70-4). (-)-JQ-1 was obtained from the SGC Oxford ((-)-JQ-1/SGCBD01, CAS: 1268524-71-5). AT1 was synthesized according to literature procedures (*23*) and detailed QC is provided in supplemental methods. Recombinant human TNF-alpha (aqueous) (210-TA-020) was purchased from R&D Systems. AZD5582 (CT-A5582, CAS: 1258392-53-8) was purchased from ChemiTek. GSK-046A (BD2-specific)(*41*) and GSK-789A (BD1-specific)(*42*) were provided by GlaxoSmithKline. Purified PHA (aqueous) (R30852801) was purchased from ThermoFisher. Recombinant IL-2 (aqueous) (200–02) was purchased from Peprotech.

#### Latency Reversal/Flow Cytometry

Cells were plated in 96-well plates at 50,000/well and treated with degraders/inhibitors for 24hrs. For 16-point titrations, cells were treated at 0, 0.001, 0.005, 0.01, 0.025, 0.05, 0.1, 0.25, 0.5, 0.75, 1, 2.5, 5, 7.5, 10, and 20 µM. For all other experiments, cells were treated at indicated concentrations. N indicates total number of biological replicates performed over three independent experiments. Cells were stained with LIVE/DEAD Fixable Aqua Dead Cell Stain (ThermoFisher) for 30min, followed by DPBS wash and fixation in 1.5% paraformaldehyde/DPBS. Cells were assayed using the iQue Screener Plus (Intellicyt) and GFP expression with dead-cell exclusion was calculated using the ForeCyt analysis software (Intellicyt). Viability calculations represent total non-aqua staining cells as a percentage of the total cell gate.

#### Western Blots

Blots were performed as previously described (*52*) with minor modification: proteins were separated by 4-20% TGX tris-glycine or 3-8% Criterion XT tris-acetate gels with tris/glycine/sds or tricine buffers respectively (BioRad) depending on size and membranes were developed using Supersignal West Atto or West Pico (ThermoFisher). Antibodies: BRD4L (Bethyl A700-005), BRD4L/S (Abcam ab128874), BRD2 (Abcam ab139690), BRD3 (ab50818), HEXIM1 (CST 12604), CDK9 (CST 2316S), CyclinT1 (CST 81464S), GAPDH (Abcam ab83956), aTubulin (Abcam ab7291), LaminB1 (Abcam ab133741), TBP (CST 44059S), Flag (Sigma F1804), V5 (Abcam ab27671).

#### Synergy

An 8-point cross titration of AZD5582 (0, 5, 10, 25, 50, 100, 250, 500 nM) and JQ1 (0, 5, 10, 25, 50, 100, 250, 500 nM), ZXH-3-26 (0, 5, 10, 25, 50, 100, 250, 500 nM), or HMBA (0, 0.1, 0.25, 0.5, 0.75, 1, 5, 10 µM) was performed in triplicate. Compounds were incubated for 24hrs and latency reversal was assayed via flow cytometry as described above. We calculated synergy based on the Bliss Independence model (*53*) like previously reported (*54, 55*) but used the maximal observed GFP induction in response to TNFα for each cell line to standardize the calculated synergy value in order to allow comparison between lines. For this work, F_a_(AZD5582, JQ1, ZXH3-26, or HMBA) = (Fraction GFP single agent – Fraction GFP DMSO)/TNFα max stimulation (JLatA2 or JLat10.6) and Fa_xy_,_O_ = (Fraction GFP combo – Fraction GFP DMSO)/ TNFα-induced GFP max (JLatA2 or JLat10.6). Graphs displaying percent GFP positive in response to combinations are not normalized and represent raw GFP values.

#### Luciferase Reporter Plasmids

We started with a previously generated pCDNA3-Luciferase construct generated in house (not published). Briefly, pCDNA3-eGFP (a gift from Doug Golenbock, Addgene plasmid #13031; http://n2t.net/addgene:13031; RRID: Addgene 13031) was digested with EcoRI and XbaI and a PCR amplified luciferase with matching restriction sites was inserted in the place of GFP, generating pCDNA3-Luci. To generate the HEXIM reporter plasmids, pCDNA3-Luci was digested with MluI/BamH1 to remove the CMV promoter. To generate pDNA3-HEXpr104-Luci, the minimal 104bp HEXIM1 promoter as reported in (*33*) was amplified from HEK293 cells along with the 716bp HEXIM1 UTR with matching restriction sites and inserted via traditional restriction cloning. To generate the control pCDNA3-HEXpr0-Luci, only the HEXIM1 UTR was amplified and cloned. Resulting constructs were then transferred to an available lentiviral vector, pLKO.1-puro (a gift from Bob Weinberg, Addgene plasmid #8453; http://n2t.net/addgene:8453; RRID: Addgene 8453) to generate stable lines. The U6 promoter was removed and MluI/XbaI sites added by deletion PCR. The HEXpr104-Luci or HEXpr0-Luci inserts digested from pCDNA3 were transferred via restriction digest cloning, generating pLKO.1-puro-HEXpr104-Luci and pLKO.1-puro-HEXpr0-Luci. At all stages, plasmids were confirmed via sanger sequencing (Genewiz/Azenta Life Sciences).

#### Luciferase Reporter Lines

Lentiviral particles were generated by transfecting 293T/17 either pLKO.1-puro-HEXpr104-Luci or pLKO.1-puro-HEXpr0-Luci, psPAX2 (a gift from Didier Trono, Addgene plasmid #12260; http://n2t.net/addgene:12260; RRID:Addgene_12260), and pMD2.G (a gift from Didier Trono, Addgene plasmid #12259; http://n2t.net/addgene:12259; RRID:Addgene_12259) using FugeneHD (Promega) per manufacturers protocol. Lentiviral particles were collected and filtered via 0.45um filter prior to transduction into fresh 293T/17 cells. Stable lines were selected using 1.5ug/mL puromycin (Selleckchem) and maintained after selection under 0.5ug/mL puromycin.

#### Luciferase Assays

Reporter lines were plated in clear bottom, white well 96-well plates and treated with compounds at indicated concentrations for 24hrs. To assess cellular viability, 10uL of the resazurin-based PrestoBlue reagent (ThermoFisher) was added to each well, incubated at 37°C for 30min, and fluorescence read using the SpectraMax M3 (Molecular Devices) using 555ex/585em with a 570nm cutoff. Immediately following the plate read, PrestoBlue containing media was removed and 100uL DMEM added to all wells. Cells were immediately lysed in Steady-Glo (Promega) per manufacturers protocol, incubated for 15min, and luminescence read on the SpectraMax M3.

#### BRD4 Overexpression Plasmids

pCDNA5-Flag-BRD4-WT (a gift from Kornelia Polyak. Addgene plasmid #90331; http://n2t.net/addgene:90331; RRID: Addgene 90331), pCDNA5-Flag-BRD4-BD (a gift from Kornelia Polyak, Addgene plasmid #90005; http://n2t.net/addgene:90005; RRID: Addgene 90005), and BRD2-GFP (a gift from Kyle Miller, Addgene plasmid #65376; http://n2t.net/addgene:65376; RRID: Addgene 65376) were purchased from Addgene. pcDNA5-Flag-BRD4-ΔCTD and pcDNA5-Flag-BRD4-BDΔCTD were generated via deletion PCR which removed amino acids 1330-1362. The pcDNA5-Flag-BRD4-CTD construct was generated via deletion PCR which removed amino acids 2-1208, leaving the 1209-1362 PID fragment previously described in (*7, 44, 45*). All BRD4 plasmids retained the n-terminal flag tag and were confirmed by sanger sequencing (Genewiz/Azenta Life Sciences). Overexpression constructs were transfected into 293T cells for 48hrs using Fugene HD (Promega) per manufacturer’s instructions. For expression analysis, cell pellets were lysed in a modified RIPA as previously described (*52*) and subject to western analysis.

#### Immunoprecipitation Experiments

For CyclinT1 IPs, Jurkat cells were treated with indicated concentrations of ZXH-3-26, JQ1, or DMSO vehicle control for 24hrs. For HEXIM IPs, 293T cells were transfected with overexpression constructs for 48hrs using Fugene HD per manufacturer’s instructions. Cell pellets were lysed in an NP-40 lysis buffer (25mM Tris pH7.5, 150nM NaCl, 1% NP-40, 1x c0mplete protease inhibitors (Millipore-Sigma), and 1X Halt Phosphatase Inhibitor cocktail (ThermoFisher) for 30min on ice. Supernatant was cleared by centrifugation and protein concentration assayed by detergent-compatible bradford (Pierce/ThermoFisher). 5uL anti-HEXIM (CST 1260) or 10uL anti-CyclinT1 (CST 81464S) were incubated with 500ug total lysate for 1hr at 4°C, followed by addition of 1mg of washed Dynabeads Protein G (ThermoFisher/Life Technologies) for an additional hour at 4°C. Beads were washed 5X in NP-40 lysis buffer and bound complexes eluted at 95°C for 10min using 50uL of western loading buffer (1X Nupage LDS buffer + 1X Nupage Reducing Agent)(ThermoFisher). Eluted IP material and input samples were subject to western analysis.

#### HIV gag RNA Induction in primary cells

Assays were performed as previously described in (*16*).

## Acknowledgments

This work was supported by CARE, a Martin Delaney Collaboratory of the National Institute of Allergy and Infectious Diseases (NIAID), the National Institute of Neurological Disorders and Stroke (NINDS), the National Institute on Drug Abuse (NIDA), and the National Institute of Mental Health (NIMH) of the National Institutes of Health, grant number 1UM1AI614567 to D.M.M., by NIAID award F30AI145588 to S.D.F., by the National Institute on Drug Abuse (NIDA) of the National Institutes of Health under award number R61DA047023/R33DA047023 to L.I.J., and by Qura Therapeutics. The iQue in the UNC Flow Cytometry Core Facility is supported in part by the Center for AIDS Research award number P30-AI050410 and the North Carolina Biotech Center Institutional Support Grant 2015-IDG-1001.

