## Supplemental Material for "BET Degraders Reveal BRD4 Disruption of 7SK and P-TEFb is Critical for Effective Reactivation of Latent HIV in CD4+ T-cells"

### Supplementary Figures

#### Supplementary Figure 1

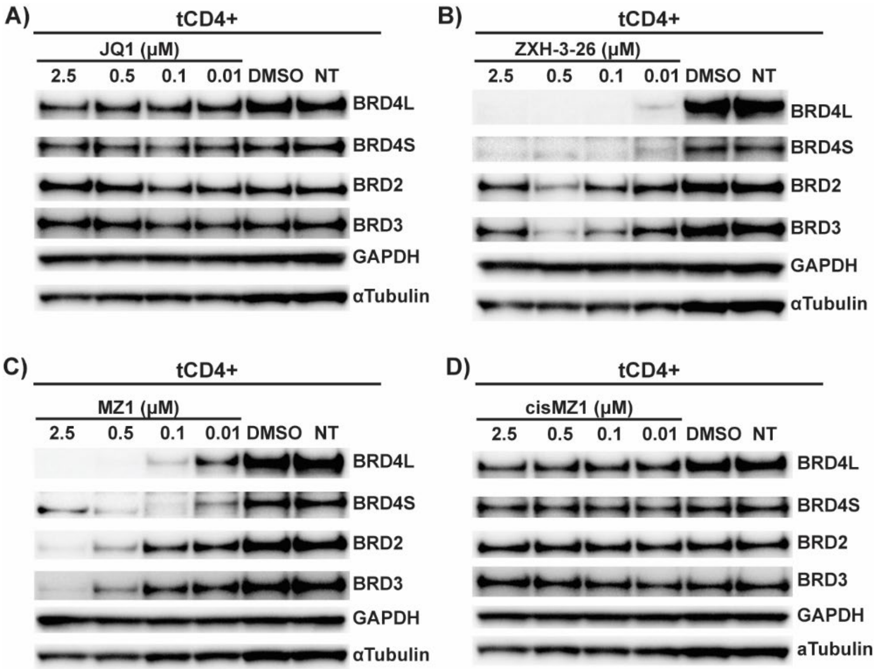


Figure S1 – *BET protein degradation in primary CD4+ T-cells* – Healthy primary human CD4+ T-cells were treated for 24 hrs at 10 nM, 100 nM, 500 nM, and 2.5 µM with (A) JQ1, (B) ZXH-3-26, (C) MZ1, and (D) control cisMZ1. BRD4L, BRD4S, BRD2, and BRD3 levels were assayed by western. GAPDH and αTubulin are provided as loading controls.

**Supplementary Figure 2**

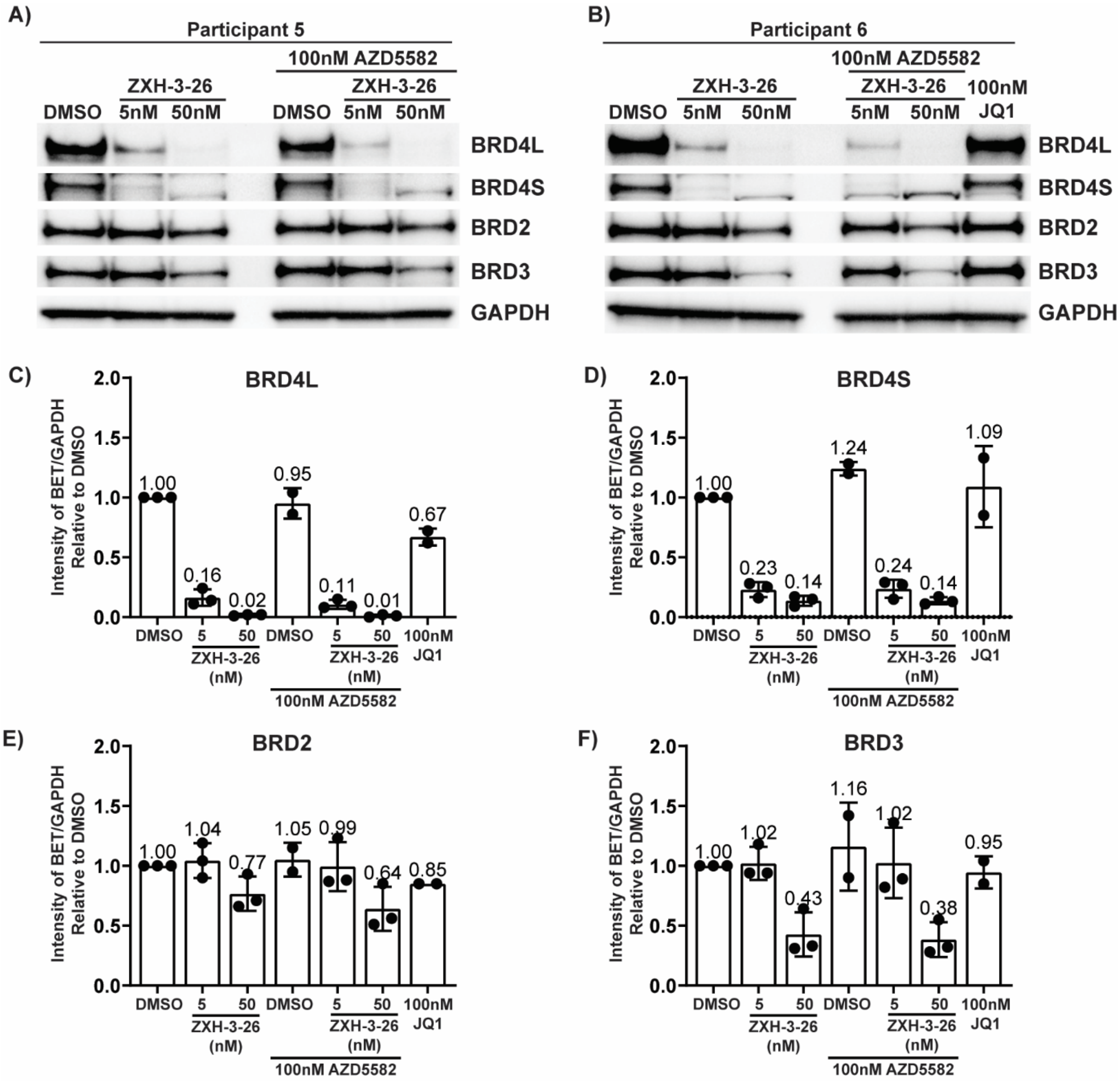


Figure S2 – *BET protein degradation by ZXH-3-26 in ART-suppressed donors* – Depending on availability of cells, 3 to 4E6 total CD4+ T-cells from ART-suppressed donors were treated with ZXH-3-26 at 5nM and 50nM, 100nM AZD5582, AZD/ ZXH-3-26, or vehicle control (DMSO). Degradation of BRD4L/S, BRD2, and BRD3 was assessed by western blot concurrent with cells treated for assessment of vRNA (*2*). Western blots for (A) participants 5 and (B) 6 are provided. The western blot for participant 7 has been previously published (*2*). Quantitation of (C) BRD4L, (D) BRD4S, (E) BRD2, and (F) BRD3 protein levels in response to ZXH-3-26 is provided for all three participants.

**Supplementary Figure 3**

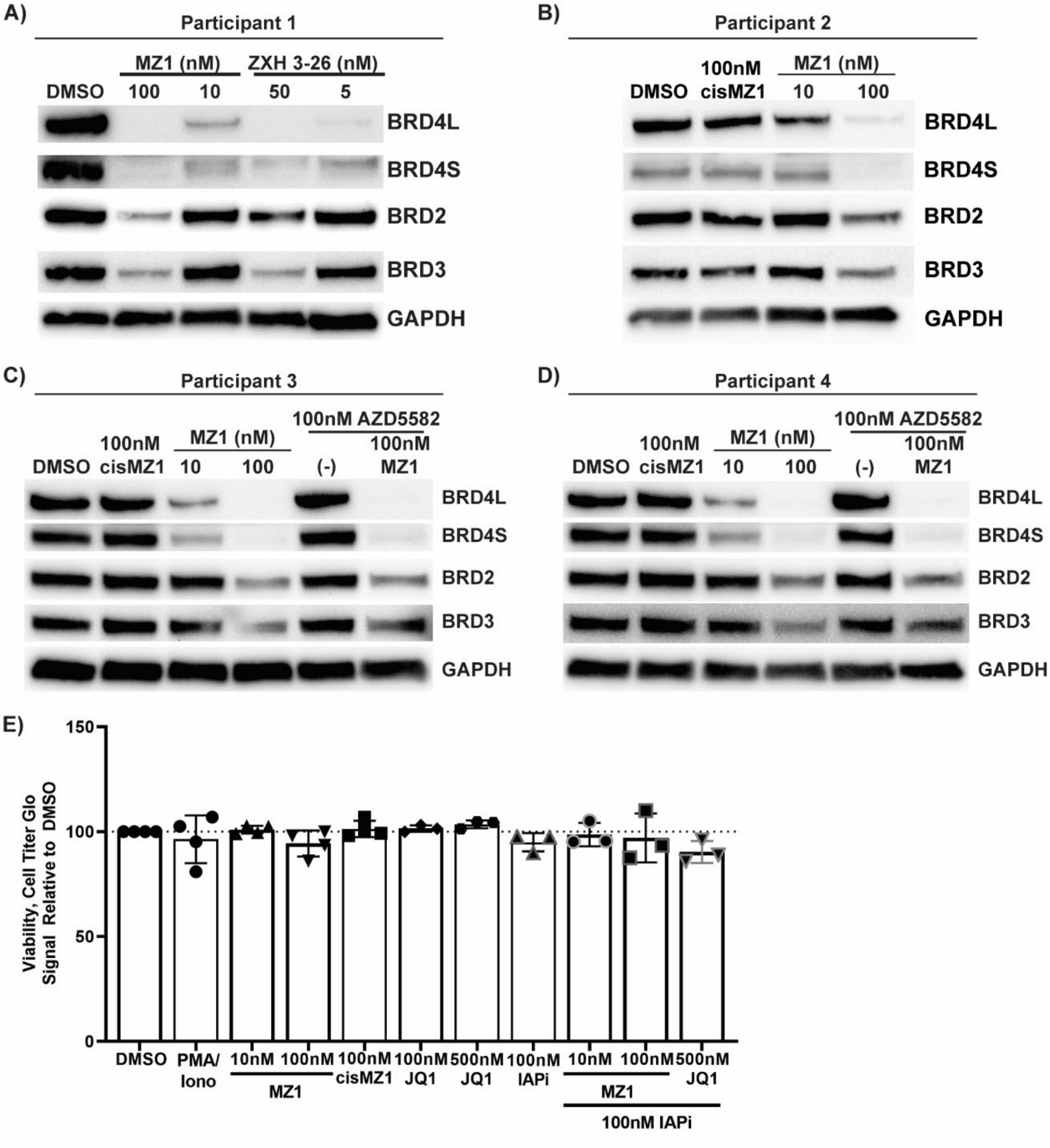


Figure S3 – *BET protein degradation by MZ1 in ART-suppressed donors* – Depending on availability of cells, 3 to 4E6 total CD4+ T-cells from ART-suppressed donors were treated with MZ1 at 10nM and 100nM, 100nM *cis*MZ1, 100nM AZD5582, AZD/MZ1, or vehicle control (DMSO) concurrent with cells treated for assessment of vRNA (Figure 4). Degradation of BRD4L/S, BRD2, and BRD3 was assessed by western blot to confirm effective on-target degradation for donors 1-4 (A-D). (E) Treated cells were assayed by CellTiter-Glo® per manufacturer's instructions to assess cytotoxicity after MZ1 treatment.

**Supplementary Figure 4**

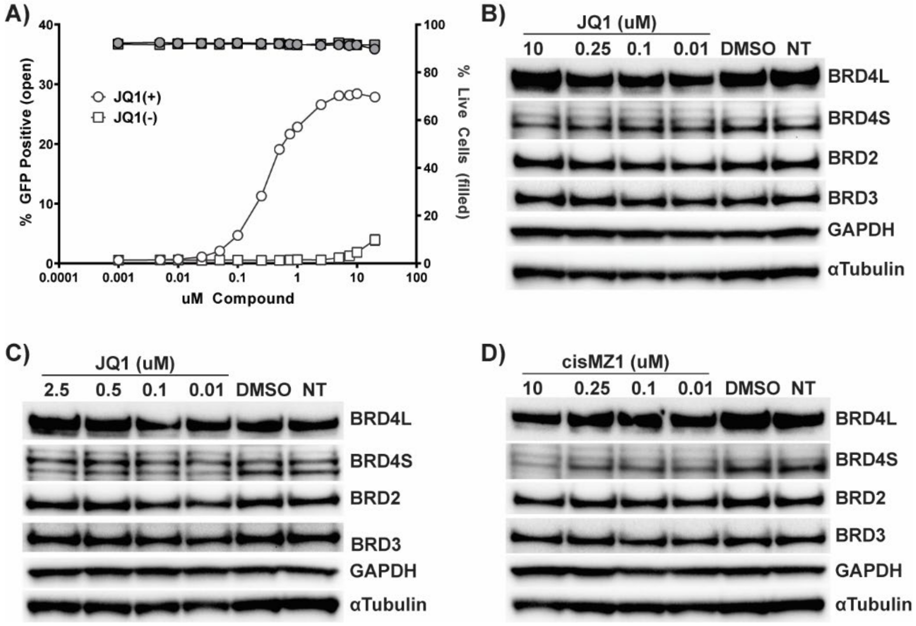


Figure S4 – *Latency reversal and targeted degradation of BET degraders in JLatA2 cells.* (A) Latency reversal by JQ1(+) versus inactive control JQ1(-) after treatment of JLatA2 cells with a 16-point dose titration for 24 hrs. Protein levels in JLatA2 cells after treatment for 24 hrs with various concentrations of JQ1 (B-C) and cisMZ1 (D).

**Supplementary Figure 5**

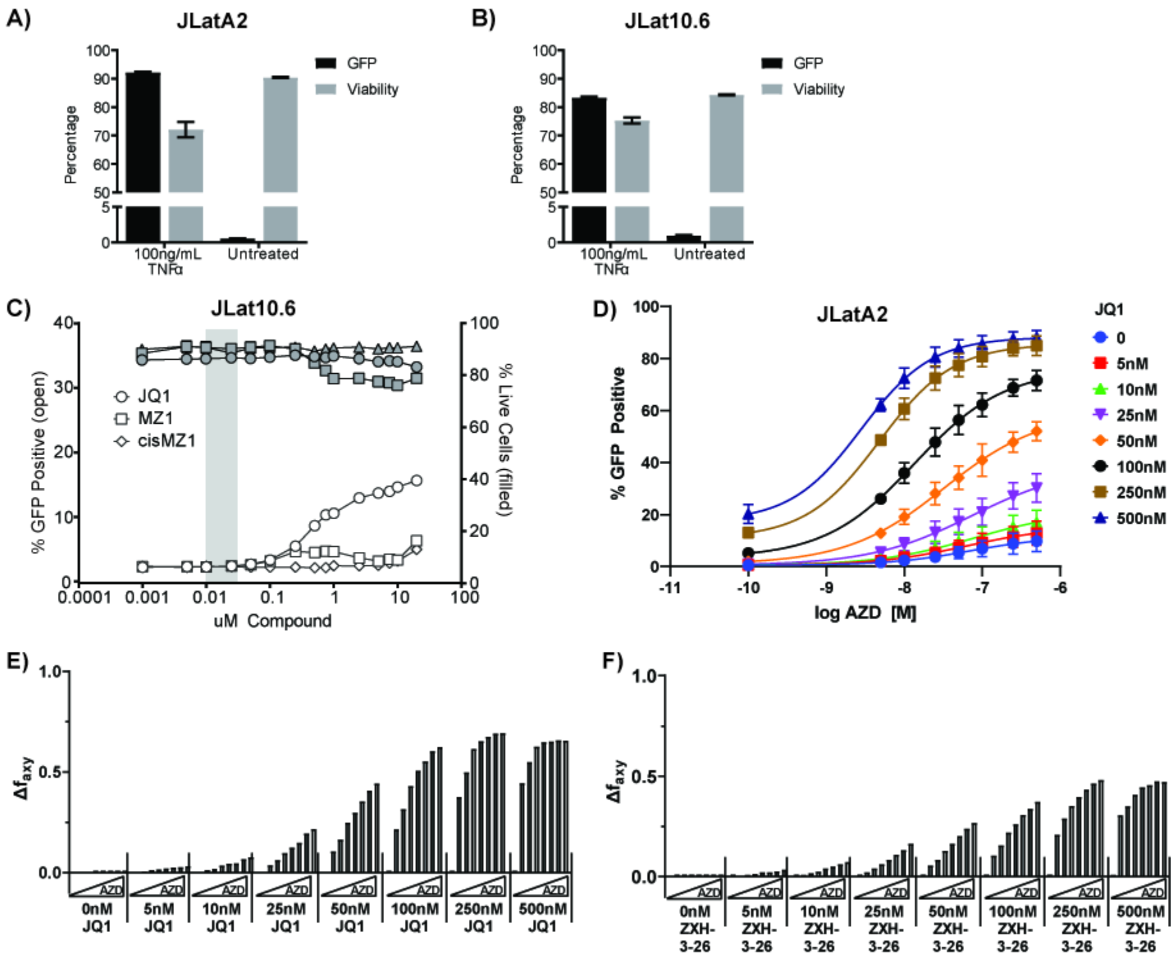


Figure S5 – *Supporting data for latency reversal and synergy in JLatA2 and JLat10.6 by IAPi/BETi/PROTACs and HMBA.* Comparison of GFP induction by 100ng/mL TNFα in (A) JLatA2 and (B) JLat10.6 cells. Latency reversal in JLat10.6 cells by (C) JQ1, MZ1 or *cis*MZ1. Each 16-point titration was performed 2 independent times with triplicate treatments for each experiment (n=6) with GFP (open symbols) assessed by flow cytometry as a measure of latency reactivation and viability by live/dead stain (gray symbols). (D) Latency reversal in JLatA2 cells with an 8-point dose titration of AZD-5582 and JQ1. Bliss synergy is observed between AZD5582 and JQ1 (E) and to a lesser extent with ZXH-3-26 (F) in JLatA2 cells.

**Supplementary Figure 6**

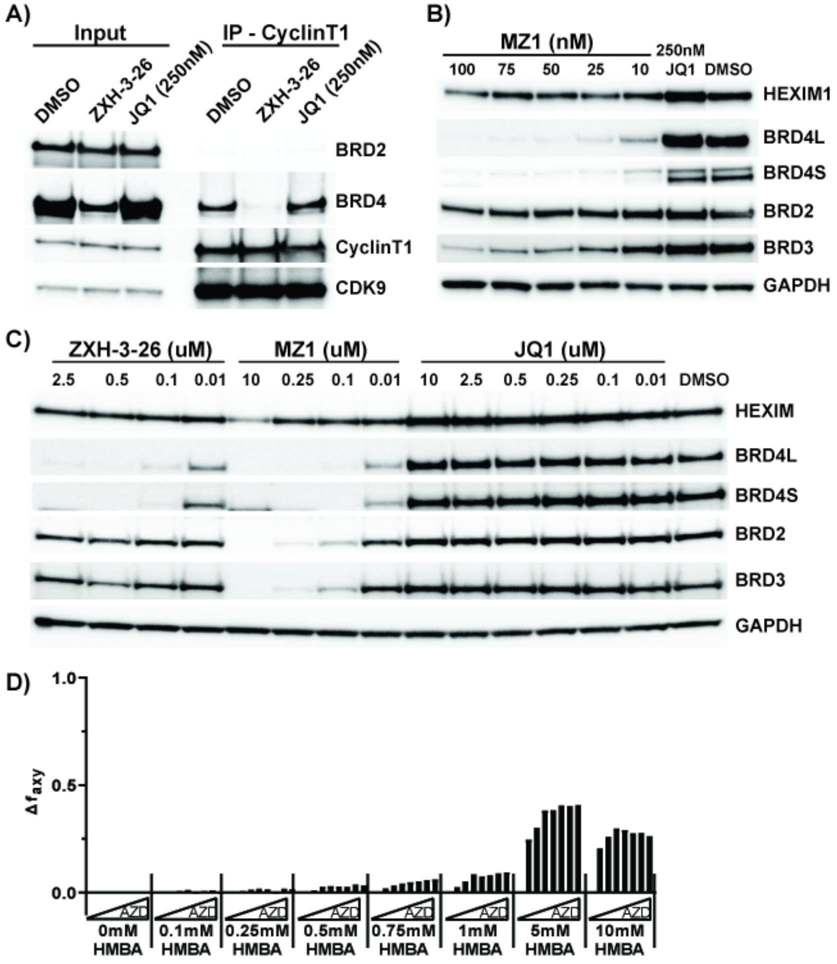


Figure S6 – *Impact of BETi or PROTACs on P-TEFb association with BRD4 and HEXIM upregulation* – (A) Jurkat cells were treated with vehicle control (DMSO), 50nM ZXH-3-26, or 250nM JQ1 for 24hrs followed by a CyclinT1 immunoprecipitation and western blot for associated proteins. HEXIM1 protein levels are not induced in (B) Jurkat-derived cells in response to MZ1. This observation is repeated in (C) extended dose curves of BET PROTACS ZXH-3-26 and MZ1 as compared to BETi JQ1. (D) Bliss synergy is observed between AZD5582 and HMBA in JLat10.6 cells.

**Supplementary Figure 7**

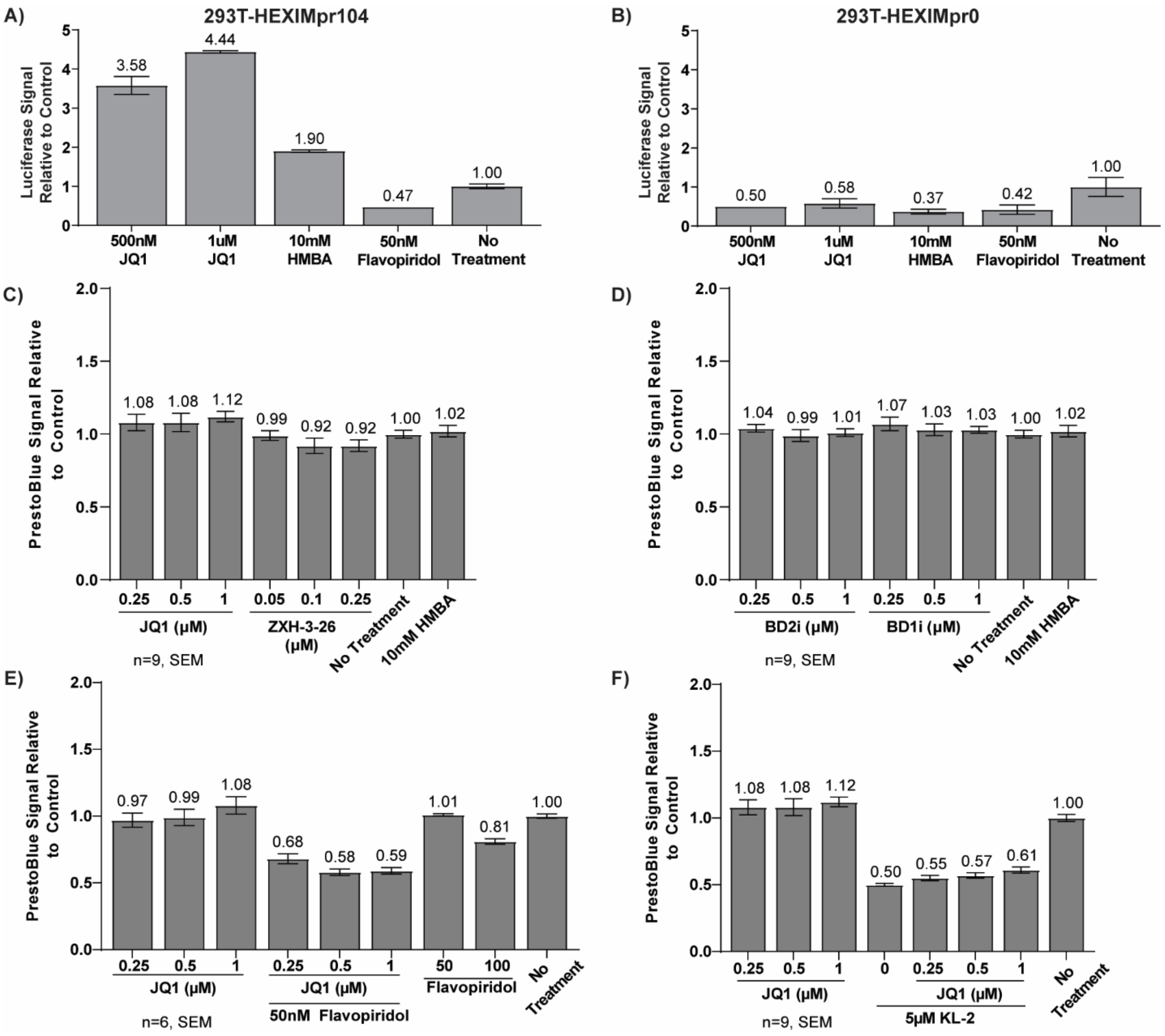


Figure S7 – *HEXIM1 Promoter Luciferase Assay* – The minimal 104bp HEXIM promoter is sufficient and responsive to known disruptors of P-TEFb JQ1 and HMBA (A) while the control reporter containing only the HEXIM1 UTR fails to induce luciferase expression (B). Viability of treated 293-HEXIMpr-Luc lines reported in Figure 7 measured by prestoblue (C-F).

**Supplementary Figure 8**

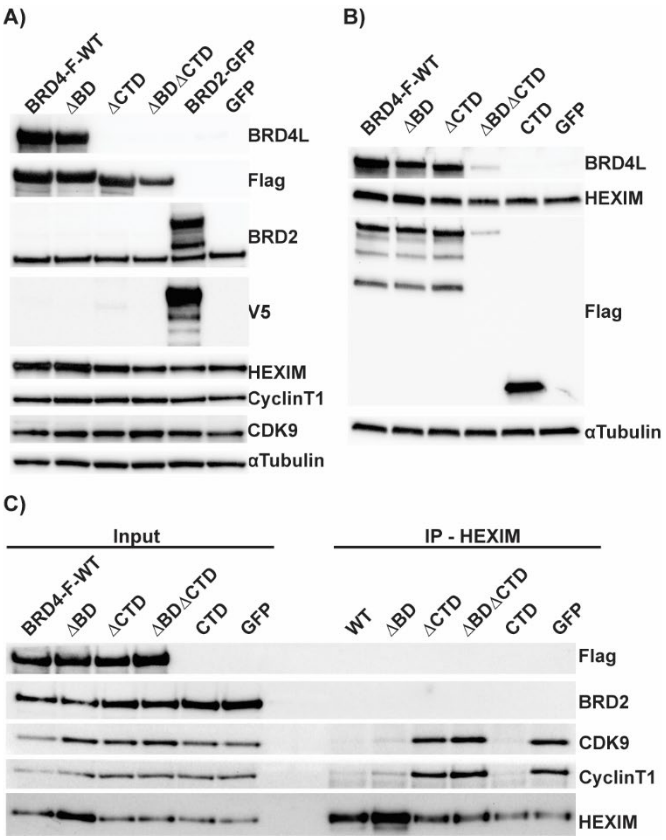


Figure S8 – *P-TEFb disruption from 7SK is dependent on the CTD but is independent of the BD domains* – Replicate experiments of (A) Overexpression plasmids containing various full-length flag-tagged BRD4 constructs or (B) full length and CTD-only constructs transfected into 293T cells for 48hrs to determine impact on HEXIM1 protein levels. (C) Overexpression constructs containing full length WT, mutant or the CTD domain alone were transfected into 293T cells for 48hrs and followed by a HEXIM1 immunoprecipitation and western blot for associated proteins.
